# Pre-operarative Multivariate Connectome Analysis for Glioma Patients

**DOI:** 10.1101/436220

**Authors:** Alessandro Crimi, Benedikt Wiestler, Jan S. Kirschke, Sandro M. Krieg

## Abstract

Recent advances in neuroimaging have allowed the use of network analysis to study the brain in a system-based approach. A system-based analysis of gliomas can shed light on mechanisms underlying neuronal connectivity and plasticity and the recovery process, and it could support surgical decision-making. Surgery has been shifting from image-guided surgery to a functional mapping-guided resection where several structural and functional modalities are used. However, reliable identification of eloquent areas during planning of surgical resection is still a challenge. Pre-operative language mapping performed by navigated transcranial magnetic stimulation (nTMS) is of great value as it elucidates functional cortical organization, which might be different from patient to patient due to the heterogeneity of the lesions and individual plasticity. In this paper we propose the construction of an “effective” speech network used for surgical decision making. This is achieved by mapping functionally relevant areas identified by nTMS on tractography-based connectomics. Subsequently we compute graph metrics on the identified speech networks of patients who show preoperative aphasia aiming to identify relevant differences between graph metrics in patients with and without preoperative aphasia. Lastly, the validity of the speech networks is examined by checking the involved graph communities.

## 1 Introduction

Glioma, which is thought to arise from glial cells, is the most common type of primary brain tumor. Gliomas are considered responsible for approximately 13000 deaths in the United States and more than 14000 deaths in Europe each year [18]. Higher-grade gliomas, in particular glioblastoma (WHO grade IV), is considered one of the most aggressive types of cancer, with a prominently invasive nature, growing along white-matter tracts. Maximum safe resection is considered a mainstay of modern glioma therapy, generally followed by chemo- and/or radiotherapy [17]. Owing to the infiltrative nature of these tumors, resection of gliomas involving relevant cortical areas is still a challenging task, and preservation of neurological function after surgery remains the goal [17]. Therefore, a deep understanding of functional and structural anatomy is crucial. Moreover, understanding of mechanisms related to plasticity, connection re-wiring, alternative and redundant routing needs to be taken into account [8]. Methods reliably visualizing neuronal connectivity are therefore highly desirable.

A connectome is a graph representation of the brain through areas (nodes) and connections. These connections can be structural, derived from tractography or functional, given by functional activations. Studies using functional connectivity observed that glioma patients have reduced connectivity in language-related areas [2] and in default mode network compared to healthy control subjects [11, 13]. Whole connectomes have also been used as discriminating signature between glioma and healthy subjects [5].

Despite advances in the new science of connectomics, techniques to directly study the function of white matter tracts *in vivo* have proved ambiguous. Cortical stimulation [7] and functional magnetic resonance imaging (fMRI) [12] are useful but constrained to identifying a focus of maximal activation. A purely anatomy-based approach such as tractography is challenging due to the definition of region of interest (ROI) and distortion of anatomical landmarks caused by lesions and brain plasticity [20]. A recent method employs navigated transcranial magnetic stimulation (nTMS) to identify seeds regions, which are followed via tractography. This has already been shown to be valuable for pre-surgical planning, both improving resection and reducing deficits in patients [14].

The arcuate fascicle is considered the most important pathway in language. It connects anterior and posterior areas, and its damage has a great impact in the language functions [19]. Nevertheless, there is a growing consensus that language is actually more distributed into large-scale cortical and subcortical networks that go beyond the arcuate fascicle, including uncinate fascicle, extreme capsule, longitudinal fascicle, and inferior fronto-occipital fascicle and related subcortical connections [6]. Moreover, glioma has traditionally been considered as a focal disease, but it is now known to in fact be a widespread, “systemic” disease across the brain [5]. Therefore, there is an increased interest from the neurological community into analysis of those networks and their pathways. A connectomic study on those pathways can shed a light on why some patients with a lesion at the same location show different deficit.

In this context we propose the definition of an “effective” or multivariate connectome, given by the fusion of functional data from nTMS and structural diffusion tensor imaging (DTI) tractography. Multivariate because it uses different modality, effective because it builds connetomes from structural data but keeping only the nodes which are truly used according to a task-based nTMS investigation. We investigated the resulting effective task-based networks in a discriminant experiment distinguishing patients presenting with pre-operative aphasia from those without aphasia in left-hemispheric tumors. The proposed method has been tested for speech tasks but it could be applied to other tasks and brain functions. The networks were compared using well known network metrics, and a novel manner to test validity of communities within the networks is also used.

## 2 Methods

### 2.1 Task-based networks construction

With the aim of combining functional and structural data, we define task-based networks. In practice, for each patient a structural connectivity matrix **A** was constructed from tractography data. Initially tractographies for all subjects have been generated by processing DTI data with the Python library Dipy [9], stemming from 2,000,000 seed-points and stopping when the fractional anisotropy was smaller than < 0.1. Tracts shorter than 30 mm or in which a sharp angle occurred have been discarded. Then the structural connectome was constructed using the 90 regions in the Automated Anatomical Labeling (AAL) atlas [21], by counting the number of tracts connecting two regions, for any pair of regions. The functional data are obtained by using nTMS. “Navigating” the brain and locating the functional areas related to a task - in our case speech - specific points in the gray matter related to the task are detected. Given the atlas registered to the volume comprising the the nTMS points, those are used as a mask leaving only the atlas ROIs where nTMS points are present. The output will be a matrix **B** constructed as follows:

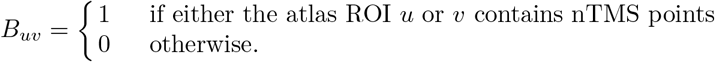

Such a matrix is used to maintain in the structural connectivity matrix **A** only the connections which are meaningful in the functional task during the nTMS, generating therefore a task-based connectivity matrix

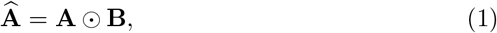

where ⨀ denotes the Hadamard or element-wise product. In our case, each 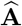 is the pre-operative speech network for each patient. These steps are depicted in Figure 1 where a resulting speech graph defined by the matrix 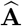 is shown together with the components needed to obtain it.

**Fig. 1.**
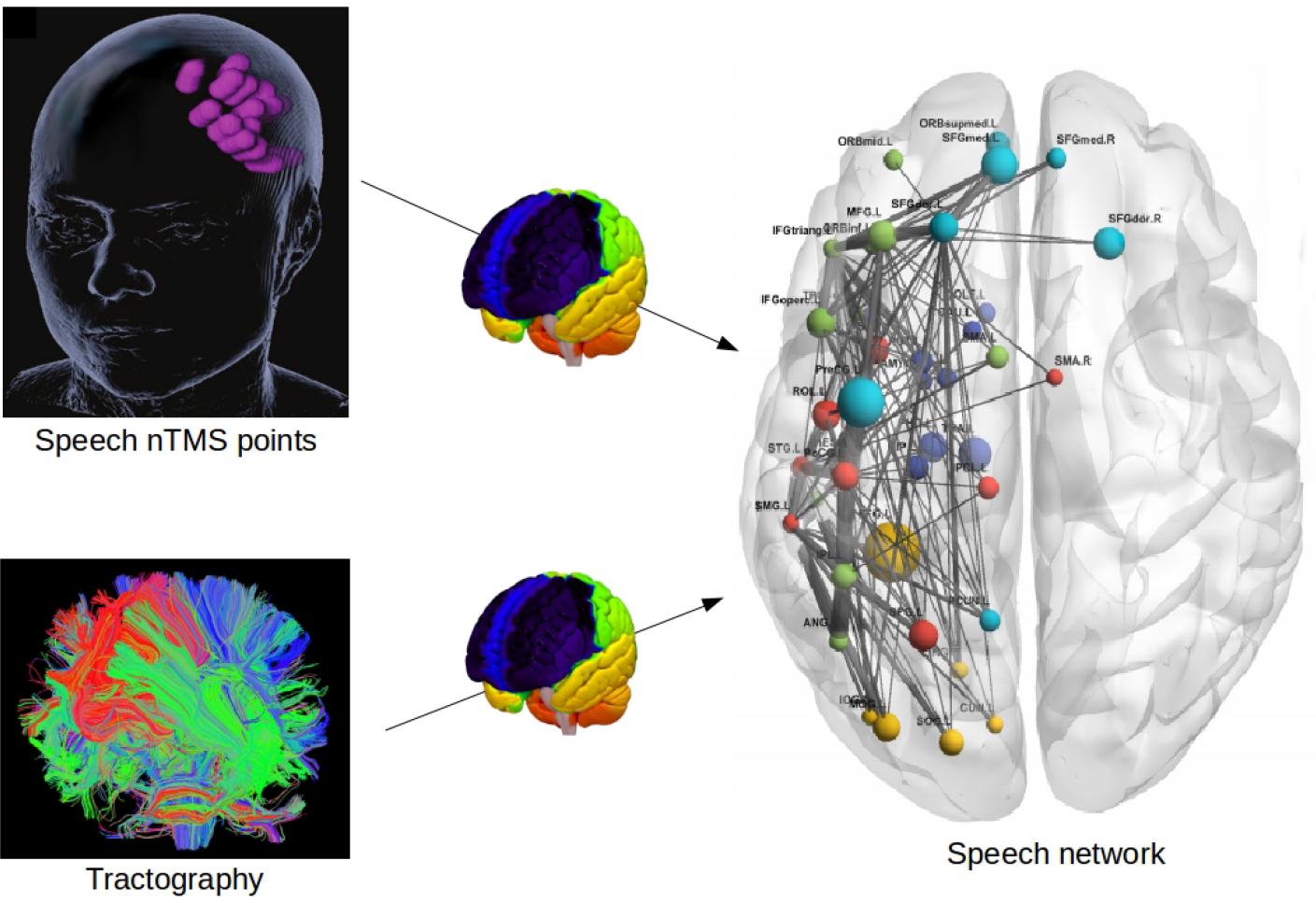
The used pipeline. Given nTMS points used to locate function and tractography, an “effective” task-based network is found.

### 2.2 Metrics

Graph metrics are important tools to analyze networks because they allow to represent the topology and efficiency of a network with only a few scalar values. Those might represent the segregation, integration, centrality, and resilience of a network [16]. Our rationale is that even in patients presenting with a similar type of tumor, individual pathways are different. To obtain this difference between sub-groups of patients graph metrics are used. Statistically significant differences are quantified by using two-tail t-tests and relative p-values. In our experiments metrics were computed at three levels:

- Globally, using the entire structural connectivity matrix **A**.
- Globally, using the speech network given by 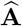.
- Locally, using the ROIs comprised in the speech network 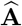.

We used only the most common metrics to be in line with previous studies on glioma [5–11, 2]. In particular, as global metrics Louvain modularity, characteristic path length, global efficiency and transitivity were used. As local metrics clustering coefficient, betweenness centrality and local efficiency were used.

The *Louvain modularity* of a network gives the degree to which the network may be subdivided into non-overlapping groups as 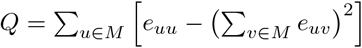 where *M* is the set of non-overlapping modules, and *e*_*uv*_ is the proportion of all links that connect nodes in the module *u* and the nodes in module *v*.

The *characteristic path length d*_*ij*_ is the average shortest path length in the network between each node *i* and *j*. The efficiency measure is given by the average inverse shortest path length. It can be computed globally or limited to the neighborhood of a node defining the local efficiency.

*Global efficiency* is defined as 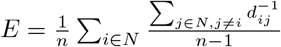.

*Transitivity* is a variation of the clustering coefficient computed directly on the global network, and it reflects the prevalence of clustered connectivity around its nodes. It can be defined as 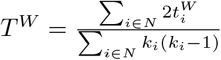, with 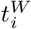 being the weighted geometric mean of triangles around node *i* and *k*_*i*_ its degree.

*Clustering coefficient* is the fraction of triangles around a node, and it can be defined as 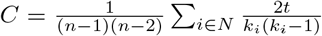 where *t*_*i*_ and *k*_*i*_ is respectively the number of triangles around a node *i*, and the degree of the node *i*.

*Betweenness centrality* is the fraction of all shortest paths in the network that passes through a given node *i* as 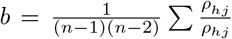 where *ρ*_*hj*_ is the number of shortest paths between *h* and *j*, and *ρ*_*hj*_ is the number of shortest paths between *h* and *j* that pass through *i*.

*Local efficiency* is the global efficiency computed on node neighborhoods.

For the local metrics, only ROIs which were common across all 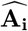 matrices were considered. A further challenge was given by the fact that 2 subjects had tumor and stimulation on the opposite hemisphere of the remaining cohort. Nevertheless, they were included and these ROIS were considered as well in the analysis even if in the opposite hemisphere.

### 2.3 Community testing

Brain functions such as language involve several brain-network communities. Despite the abundance of contributions related to community detection there is no widespread consensus defining what communities are and how to test their significance. This has emerged as a need since recent neuroscience studies suggests that conscience and brain functions emerge from community interactions, but care should be made on existing tools for community detection [1].

Using Markov random walk, Piccardi introduced the persistence probability as a measure of the strength of a community in a graph with salient community structure [15]. From an adjacency matrix **A** an N-state Markov chain can be defined by a transition matrix **P** by performing a row-normalization on **A**, as 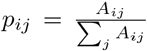, expressing the transition probability from node *i* and *j*. Then, it is possible to define a K-state Markov chain with transition probability as 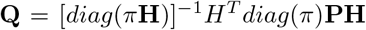, where **H** is the matrix coding the partition obtained by Louvain modularity or other methods, and *π* is defining the rule *π* = *π***P** which represents the equilibrium distribution of a random walk [15]. The diagonal of the matrix 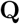 gives the persistence probability, which is the level of significance of the partitioning. Therefore we compute the persistence probability of the partitioning detected by the Louvain modularity described in the previous section to see if there is a difference within the communities detected for the sub-groups and towards a random scale free network with the same nodes of the used atlas.

## 3 Data and Experimental Settings

The cohort comprised 17 patients from the TUM University Hospital, being in average 57.3 α 8.8 years old, and with tumor of different grades and locations. All patients underwent MRI scanning before surgery, and pre-operative aphasia was evaluated looking for speech and impairment of verbal communication according to an established grading [19]. In this manner the cohort was subdivided into subjects with aphasia (WA) n=8, and subjects without aphasia (NA) n=9. DTI, T1-weighted volumes and positions of nTMS were acquired and co-registered onto an atlas by using an affine linear registration with 12 degrees of freedom. The diffusion brain volumes were acquired with a Philips Achieva 3 Tesla with the following imaging parameters: TR = 8500 ms, TE = 60 ms, flip angle 90° using 32 gradient directions. The T1-weighted brain volumes were acquired with the following imaging parameters: TR = 9 ms, TE = 4 ms, and 1 *mm*^3^ isovoxel covering the whole head. Performing an object-namimg task according to the most recent protocol was used for nTMS [14], the magnetic stimulation coil was manually moved in steps of approximately 10 mm covering most of the hemisphere where the lesion was present. The coordinates of relevant points for the speech task were exported through the Nexstim eXimia software (Nexstim Oy, Helsinki, Finland).

## 4 Results and Discussions

None of the global metrics applied to the whole brain were significant in discriminating the two groups of patients, with (WA) and without (NA) pre-operative aphasia.

The results of the network analysis for the speech networks are reported in 1 and for the common areas are reported in Table 2 and 3. Given the heterogeneity of lesions and the individual mapping only two regions resulted common across all subjects: The inferior parietal gyrus and superior parietal gyrus. Those are regions in the brain known to be related to speech processing and understanding [3].

**Table 1.** Global graph metrics for the speech network.

| Features | WA sub-group | NA sub-group | p-value |
| --- | --- | --- | --- |
| Louvain modularity | 0.518 ($\pm 0.020$) | 0.511 ($\pm 0.015$) | 0.444 |
| Char. path length | 2.787 ($\pm 0.215$) | 0.222 ($\pm 0.123$) | 0.041 |
| Global efficiency | 0.334 ($\pm 0.040$) | 0.375 ($\pm 0.025$) | 0.056 |
| Transitivity | 0.013 ($\pm 0.002$) | 0.012 ($\pm 0.002$) | 0.990 |

**Table 2.**
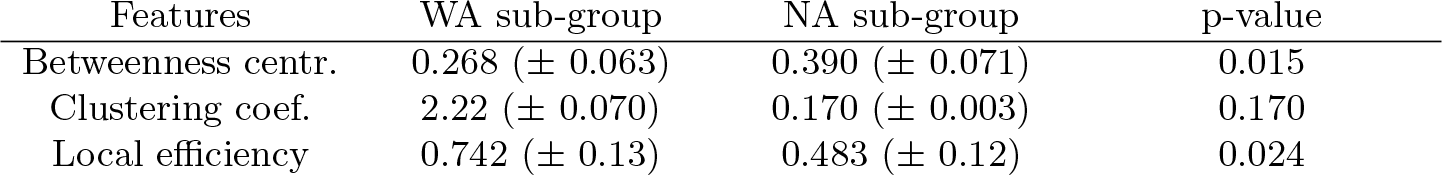
Local graph metrics for the inferior parietal gyrus across the dataset.

| Features | WA sub-group | NA sub-group | p-value |
| --- | --- | --- | --- |
| Betweenness centr. | 0.268 ($\pm 0.063$) | 0.390 ($\pm 0.071$) | 0.015 |
| Clustering coef. | 2.22 ($\pm 0.070$) | 0.170 ($\pm 0.003$) | 0.170 |
| Local efficiency | 0.742 ($\pm 0.13$) | 0.483 ($\pm 0.12$) | 0.024 |

**Table 3.**
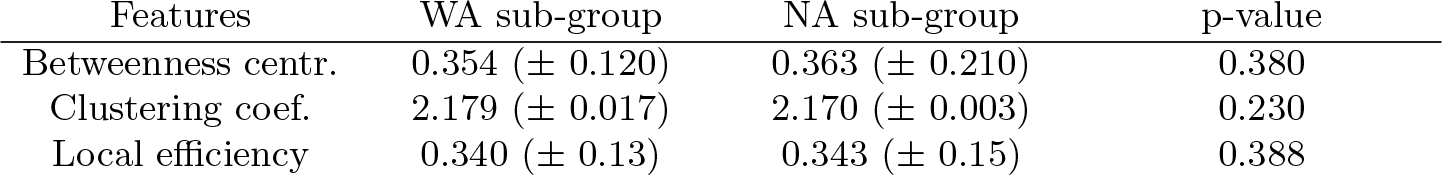
Local graph metrics for the superior parietal gyrus across the dataset.

| Features | WA sub-group | NA sub-group | p-value |
| --- | --- | --- | --- |
| Betweenness centr. | 0.354 (± 0.120) | 0.363 (± 0.210) | 0.380 |
| Clustering coef. | 2.179 (± 0.017) | 2.170 (± 0.003) | 0.230 |
| Local efficiency | 0.340 (± 0.13) | 0.343 (± 0.15) | 0.388 |

The processes supporting speech production and comprehension within the brain take advantage of widespread networks and region. Tumor lesions can destroy gray matter region that serves as a node of the network ultimately disrupting certain communication within the brain. During the analysis of the speech networks, it was noted that the characteristic path length and global efficiency were statistically significantly different between the two sub-groups. In particular the characteristic path length was larger for the group with aphasia implying that given the neuronal pathways to relate local communities or distant regions are more cumbersome or suggesting the lack of redundancy and plasticity ultimately leading to aphasia. Lastly, the communities detected by using the Louvain modularity showed similar persistence probabilities comparing the two sub-groups but superior to the persistence on the random scale free network. Namely, the mean *q* values for the communities of the WA sub-group, NA sub-group and random network were respectively 0.652, 0.631 and 0.3, showing that detected have higher probability of being valid compare to random scale free networks.

The AAL atlas is known to have limitations as a rather coarse parcellation [10]. However, initial experiments conducted with more refined atlas were not satisfactory after visual inspection due to failed registrations even using more advanced non-rigid methods. Registration of brain with large lesions to atlas is still an open issue given the heterogeneity of shape and location of lesions which cannot comprehensively be managed by existing methods [4]. Therefore we preferred to use the AAL which despite the coarse parcellation was properly registered to the data even using standard linear registration. Future works comprises the visualization and navigation of the networks detected with the proposed approach.

## 5 Conclusions

A method to map functional detected areas on a connectome is proposed and the validity of the produced networks are analyzed. Speech production and comprehension requires a complex, and plastic network within the brain, and it is therefore the application used for the approach. Although larger studies are needed, the comparison between detected speech networks of patients presenting aphasia and those without were significant while a whole-brain metrics comparison was not significant, proving the usefulness of the method during surgical planning and potentially to predict eventual post-operative deficits.

## References

1. Bassett, D.S., Porter, M.A., Wymbs, N.F., Grafton, S.T., Carlson, J.M., Mucha, P.J.: Robust detection of dynamic community structure in networks. Chaos: An Interdisciplinary Journal of Nonlinear Science 23(1), 013142 (2013)

2. Briganti, C., Sestieri, C., Mattei, P., Esposito, R., Galzio, R., Tartaro, A., Romani, G., Caulo, M.: Reorganization of functional connectivity of the language network in patients with brain gliomas. American Journal of Neuroradiology 33(10), 1983–1990 (2012)

3. Buchsbaum, B.R., Hickok, G., Humphries, C.: Role of left posterior superior temporal gyrus in phonological processing for speech perception and production. Cognitive Science 25(5), 663–678 (2001)

4. Cuadra, M.B., Pollo, C., Bardera, A., Cuisenaire, O., Villemure, J.G., Thiran, J.P.: Atlas-based segmentation of pathological mr brain images using a model of lesion growth. IEEE transactions on medical imaging 23(10), 1301–1314 (2004)

5. Derks, J., Dirkson, A.R., de Witt Hamer, P.C., van Geest, Q., Hulst, H.E., Barkhof, F., Pouwels, P.J., Geurts, J.J., Reijneveld, J.C., Douw, L.: Connectomic profile and clinical phenotype in newly diagnosed glioma patients. NeuroImage: Clinical 14, 87–96 (2017)

6. Dick, A.S., Tremblay, P.: Beyond the arcuate fasciculus: consensus and controversy in the connectional anatomy of language. Brain 135(12), 3529–3550 (2012)

7. Duffau, H.: Stimulation mapping of white matter tracts to study brain functional connectivity. Nature Reviews Neurology 11(5), 255 (2015)

8. Fornito, A., Zalesky, A., Breakspear, M.: The connectomics of brain disorders. Nature Reviews Neuroscience 16(3), 159 (2015)

9. Garyfallidis, E., Brett, M., Amirbekian, B., Rokem, A., Van Der Walt, S., Descoteaux, M., Nimmo-Smith, I., Contributors, D.: Dipy, a library for the analysis of diffusion mri data. Frontiers in neuroinformatics 8 (2014)

10. Gordon, E.M., Laumann, T.O., Adeyemo, B., Huckins, J.F., Kelley, W.M., Petersen, S.E.: Generation and evaluation of a cortical area parcellation from resting-state correlations. Cerebral cortex 26(1), 288–303 (2014)

11. Harris, R.J., Bookheimer, S.Y., Cloughesy, T.F., Kim, H.J., Pope, W.B., Lai, A., Nghiemphu, P.L., Liau, L.M., Ellingson, B.M.: Altered functional connectivity of the default mode network in diffuse gliomas measured with pseudo-resting state fmri. Journal of neuro-oncology 116(2), 373–379 (2014)

12. Hart, M.G., Price, S.J., Suckling, J.: Connectome analysis for pre-operative brain mapping in neurosurgery. British journal of neurosurgery 30(5), 506–517 (2016)

13. Kinno, R., Ohta, S., Muragaki, Y., Maruyama, T., Sakai, K.L.: Differential reorganization of three syntax-related networks induced by a left frontal glioma. Brain 137(4), 1193–1212 (2014)

14. Krieg, S.M., Sollmann, N., Obermueller, T., Sabih, J., Bulubas, L., Negwer, C., Moser, T., Droese, D., Boeckh-Behrens, T., Ringel, F., et al.: Changing the clinical course of glioma patients by preoperative motor mapping with navigated transcranial magnetic brain stimulation. BMC cancer 15(1), 231 (2015)

15. Piccardi, C.: Finding and testing network communities by lumped markov chains. PloS one 6(11), e27028 (2011)

16. Rubinov, M., Sporns, O.: Complex network measures of brain connectivity: uses and interpretations. Neuroimage 52(3), 1059–1069 (2010)

17. Sanai, N., Berger, M.S.: Glioma extent of resection and its impact on patient outcome. Neurosurgery 62(4), 753–766 (2008)

18. Siegel, R.L., Miller, K.D., Jemal, A.: Cancer statistics, 2016. CA: a cancer journal for clinicians 66(1), 7–30 (2016)

19. Sollmann, N., Ille, S., Hauck, T., Maurer, S., Negwer, C., Zimmer, C., Ringel, F., Meyer, B., Krieg, S.M.: The impact of preoperative language mapping by repetitive navigated transcranial magnetic stimulation on the clinical course of brain tumor patients. BMC cancer 15(1), 261 (2015)

20. Southwell, D.G., Hervey-Jumper, S.L., Perry, D.W., Berger, M.S.: Intraoperative mapping during repeat awake craniotomy reveals the functional plasticity of adult cortex. Journal of neurosurgery 124(5), 1460–1469 (2016)

21. Tzourio-Mazoyer, N., et al.: Automated anatomical labeling of activations in SPM using a macroscopic anatomical parcellation of the MNI MRI single-subject brain. Neuroimage 15(1), 273–289 (2002)

